# DETECTION OF BIOSYNTHETIC GENE CLUSTERS FROM METAGENOME OF NATURAL HONEY COLLECTED IN VIETNAM

**DOI:** 10.64898/2026.04.03.716293

**Authors:** Hoang-Nam Nguyen, Oanh Thi Phuong Kim, Thuy Thi Tran

## Abstract

The antimicrobial properties of natural honey are partly attributed to bioactive secondary metabolites produced by its associated microbial communities, yet the biosynthetic capacity of these communities remains poorly characterized. Here, we applied a metagenomic approach to investigate the biosynthetic gene cluster (BGC) diversity of bacteria associated with *Apis cerana* honey from the Northwest mountainous region of Vietnam – a biogeographically distinct and underexplored ecosystem. A total of 366 BGCs spanning 38 compound classes were identified, with terpenes, nonribosomal peptide synthetases (NRPS), and ribosomally synthesized and post-translationally modified peptides (RiPPs) being the most prevalent. Strikingly, 304 BGCs (>83%) lacked close matches in the MIBiG reference database, indicating a high degree of biosynthetic novelty relative to previously characterized natural product repertoires. Among the identified clusters, an azole-containing RiPP BGC recovered from a metagenome-assembled genome (MAG) assigned to *Atlantibacter hermannii* was predicted to exhibit strong antibacterial activity, with a probability score of 74.5%, representing a prioritized target for heterologous expression and bioactivity validation. These findings establish the *Apis cerana* honey microbiome as a tractable and largely untapped reservoir for applied microbial research, with direct implications for the discovery of novel antimicrobial agents from underexplored environmental niches.

**Importance:** Antimicrobial resistance represents one of the most pressing challenges in modern medicine, driving urgent demand for novel bioactive compounds from underexplored microbial sources. Honey-associated bacterial communities are recognized contributors to the antimicrobial properties of natural honey, yet their biosynthetic capacity remains poorly characterized at the metagenomic level. This study demonstrates that the microbiome of *Apis cerana* honey from a biogeographically distinct region of Vietnam harbors extensive and largely novel biosynthetic gene cluster diversity, with over 83% (304/366) of identified BGCs lacking database references. These findings position honey microbiomes as a tractable and underutilized reservoir for applied microbial research, with direct relevance to the discovery of new antimicrobial agents. The identification of a candidate antibacterial RiPP BGC in *Atlantibacter hermannii* provides a concrete target for future cultivation-based and heterologous expression studies, bridging metagenomic discovery with applied biotechnological pipelines.

## Introduction

Honey has been used for thousands of years for its medicinal properties, one of which is its antimicrobial activity. The antimicrobial activity of honey has been demonstrated to be effective against a wide variety of pathogens, including antibiotic-resistant strains (Combarros-Fuertes et al. 2020). As a traditional remedy, honey is occasionally used as a natural alternative to antibiotics in the treatment of certain infections, such as wound healing and upper respiratory tract infections (Al-Waili 2004; Abuelgasim et al. 2021). Various antimicrobial compounds in honey, which combine to provide broad-spectrum antimicrobial activity, are produced by microorganisms. The release of these compounds is inherently in competition with other microbes for limited resources such as nutrients. These compounds can kill or inhibit the growth of other microorganisms in their environment, giving the producing microbe a competitive advantage. Honey microflora is considered a potential source of antimicrobial compounds with applications in the food and agrochemical industries (Lee et al. 2008).

The development of next-generation sequencing technology and bioinformatic tools has revolutionized the "omics" research field, in which metagenomics has emerged as a powerful method for characterizing environmental DNA (eDNA). The method enables researchers to gather and examine information from the genomes of all living things in an environmental sample (Hugenholtz and Tyson 2008). Traditional microbiological methods are based on cultivating microbial species in a laboratory, which can be time-consuming and biased towards only culturable microorganisms. On the other hand, metagenomics enables the direct sequencing and characterization of eDNA, including the DNA of microorganisms that cannot be cultured (Hugenholtz and Tyson 2008; Culligan et al. 2014). This provides a significantly more extensive understanding of the diversity and the function of microbial communities. Their genomes can also be recovered from metagenome by assembly and binning tools. The potential to screen for new gene clusters has been made possible by the recovery of metagenome-assembled genomes (MAGs) from these uncultivable organisms (Parks et al. 2017). It has unlocked the metabolic potential of previously underutilized metabolic processes, such as secondary metabolism gene clusters.

Secondary metabolites are produced by microorganisms in the environment but are not directly related to the growth of these microorganisms. These compounds possess various physiological functions that enable the host producers to thrive in particular environments. Microorganisms tend to organize their genes involved in the synthesis of secondary metabolites into clusters known as biosynthetic gene clusters (BGCs) (Medema et al. 2015; Chen et al. 2020). They are gene clusters that encode a biosynthetic pathway for producing secondary metabolites and their chemical variants, including antimicrobial compounds. The biosynthesis of these compounds is typically regulated by a complex network of genes, enzymes, and signaling pathways (Medema et al. 2015). Since the multidrug resistance of pathogens is becoming a pressing global problem (Chinemerem Nwobodo et al. 2022), it is essential to look for alternative antimicrobial agents. The identification and characterization of BGCs in microorganisms are significant for the discovery of antimicrobial drugs; for example, 13 complete BGCs, encoding type II polyketides, were identified from 3,203 metagenomic samples of the human microbiome with a notable presence in the gut, mouth, and skin, and evidence of transcription under host-colonization conditions. The experimental characterization of two selected BGCs led to the successful purification and structural determination of five new type II polyketide molecules, two of which exhibited potent antibacterial activity against competing microbiome members (Sugimoto et al. 2019). In Antarctic soil, over 1,400 full-length BGCs were recovered from uncultivated bacteria using long-read sequencing and genome mining, including a potential novel family of RiPPs in the Gammaproteobacterial order UBA7966, highlighting the rich BGC diversity in underexplored phyla and the potential of long-read sequencing for accessing novel antimicrobial compounds (Waschulin et al. 2022).

Although the antimicrobial, anti-oxidative compounds in the honey may come from plant origin (such as phenolic acids and flavonoids) (McLoone et al. 2016) or bees (like bee-defensin 1) (Proaño et al. 2021) have been described in detail, the compounds are synthesized by microorganisms in honey has not yet received much attention. In particular, to date, there have been no studies exploring the BGCs of the honey microbiome. In this study, we used a sequence-based metagenomic approach to recover bacterial MAGs and investigate the BGCs of the recovered genomes in raw natural honey.

## Materials and Methods

### Metagenomic data

In this study, raw reads of metagenomic sequence datasets from nine honey samples collected in the Northwest region of Vietnam were used for analysis. The datasets from these honey samples were described previously and available in the NCBI Sequence Read Archive (SRA) with the accession numbers SRR23325837, SRR23325836, SRR23325835 (Nguyen et al. 2025a) and SRR26265549, SRR26265548, SRR2626555, SRR26265550, SRR26265547, SRR26265546 (Nguyen et al. 2025b). Taxonomic classification of sequencing reads was determined by Kraken2 (Wood et al. 2019), after which bacterial reads together with unclassified reads of all nine honey samples were combined for contig co-assembly using MEGAHIT (v1.2.9) (Li et al. 2015). Only contigs with a minimum length of 500 bp were retained.

### Reconstruction of MAGs

The assembled contigs were used to reconstruct genomes. Firstly, the reads were mapped back to the assembled contigs using BBmap (v39.01) (Bushnell 2014) to generate the coverage information. These information were used by the software tools metaBAT2 (v2.15) (Kang et al. 2019), SemiBin2 (v2.2.0) (Pan et al. 2023), and Vamb (v5.0.4) (Nissen et al. 2021) to group contigs into genome bins using default parameters. SemiBin2 and Vamb utilized only contigs with a minimum length of 1,000 bp, whereas metaBAT2 used only contigs with a minimum length of 1,500 bp as input data. The bins generated by the three tools were subsequently refined using DAS Tool (Sieber et al. 2018) to produce an optimized bin set. Bin quality was assessed using CheckM2 (v1.1.0) (Chklovski et al. 2023). Bins with completeness ≥ 50% and contamination ≤ 10% were considered metagenome-assembled genomes (MAGs). Contigs ≥ 5,000 bp which were not assigned to MAGs (namely “unbinned_5000.fa” ) were subsequently analyzed together with the MAGs.

Taxonomic assignment of MAGs was performed using GTDB-Tk (v2.4.1) (Chaumeil et al. 2020) with at least 70% of marker genes were covered and the Bin Annotation Tool (v6.0.1) (Von Meijenfeldt et al. 2019) with at least 50% of the bit-score evidence, both employing the GTDB database release R220 (Chaumeil et al. 2022). The classification results obtained from the two tools were cross-compared to ensure reliability. Contigs containing sequences of interest (biosynthetic gene clusters) that were not assigned to MAGs were taxonomically classified using the Contig Annotation Tool (v6.0.1) (Von Meijenfeldt et al. 2019) with at least 50% of the bit-score evidence and the GTDB database release R220 (Chaumeil et al. 2022).

### Analysis of secondary metabolism gene clusters

The MAGs and the “unbinned_5000.fa” file were analyzed using antiSMASH (v8.0.2) (Blin et al. 2025) to detect biosynthetic gene clusters (BGCs) involved in the production of secondary metabolites. For BGC quantification, individual biosynthetic components within hybrid BGC sequences were counted separately. For example, a hybrid BGC annotated as “NRPS + T1PKS” was counted as one NRPS-class BGC and one T1PKS-class (type I polyketide synthase) BGC.

A similarity network of homologous BGC sequences was constructed using BiG-SCAPE (v2.0) (Navarro-Muñoz et al. 2020) with the latest version of database MIBiG database 4.0 (Zdouc et al. 2025) and 0.3 cut-off BGC similarity value and was subsequently visualized using Cytoscape (v3.9.1) (Shannon et al. 2003). BGC sequences belonging to the same gene cluster family (GCF) were compared and visualized using clinker (v0.0.31) (Gilchrist and Chooi 2021).

The GenBank files containing BGC information generated by antiSMASH were used as input data to predict the properties of the corresponding secondary metabolites using NPBdetect (v1.1.0) (Goyat et al. 2025).

The functions of genes within BGCs of interest were determined through amino acid sequence similarity searches using BLASTp (web-based) against the *RefSeq protein* database and the NCBI *non-redundant* (*nr*) database (Johnson et al. 2008). In addition, NeuRiPP (De Los Santos 2019) and RiPPMiner (Agrawal et al. 2017) were employed to screen peptide sequences for their potential as precursor peptides of RiPP-class secondary metabolites.

## Results

### Recovery of MAGs and taxonomic classification

Using DAS Tool, an optimized bin set comprising 65 bins was selected from three independent bin sets reconstructed by metaBAT2, SemiBin2, and Vamb. Quality assessment with CheckM2 indicated that 42 bins met the quality criteria, with completeness ≥ 50% and contamination ≤ 10% (**Table 1**).

**Table 1.**
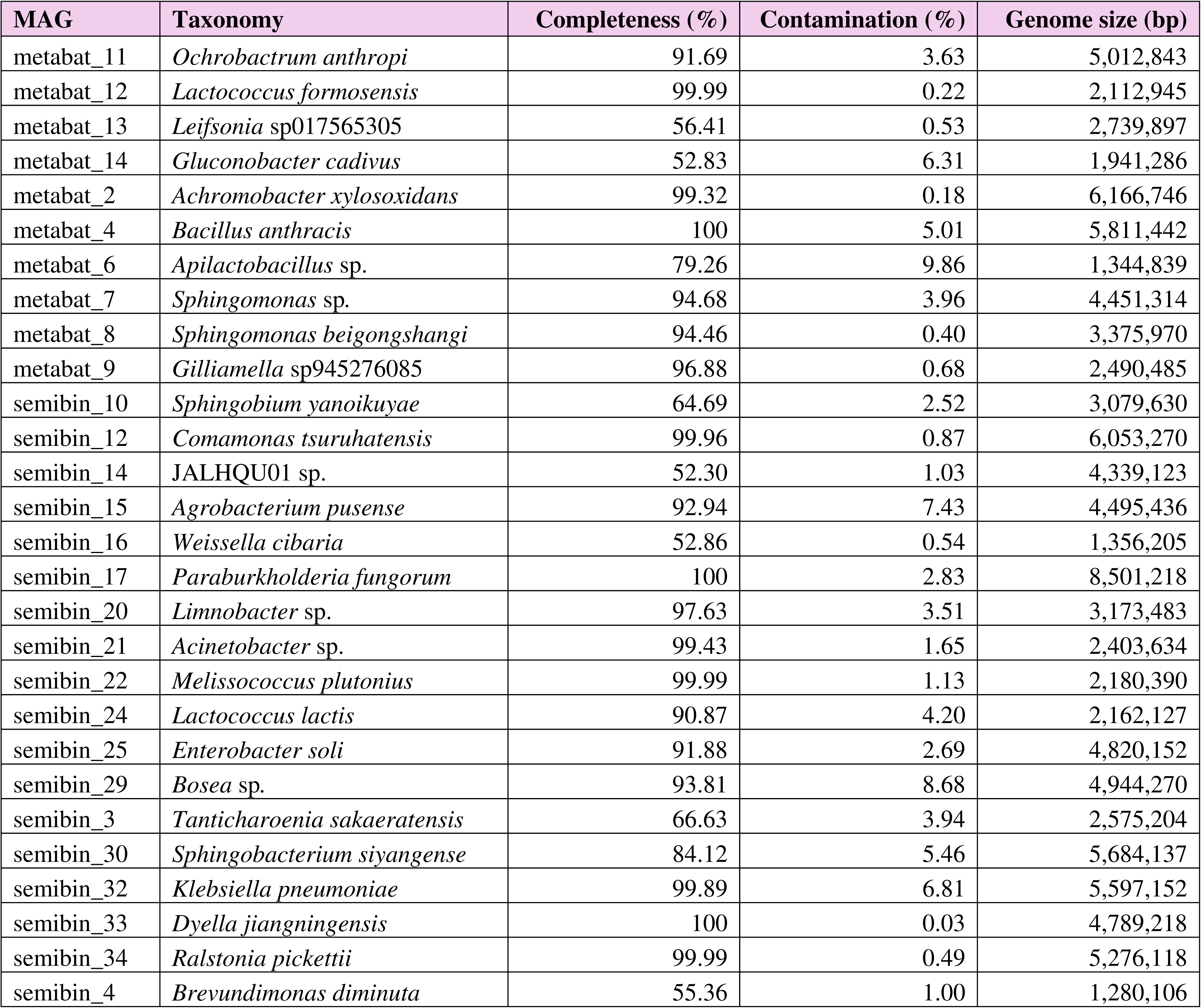
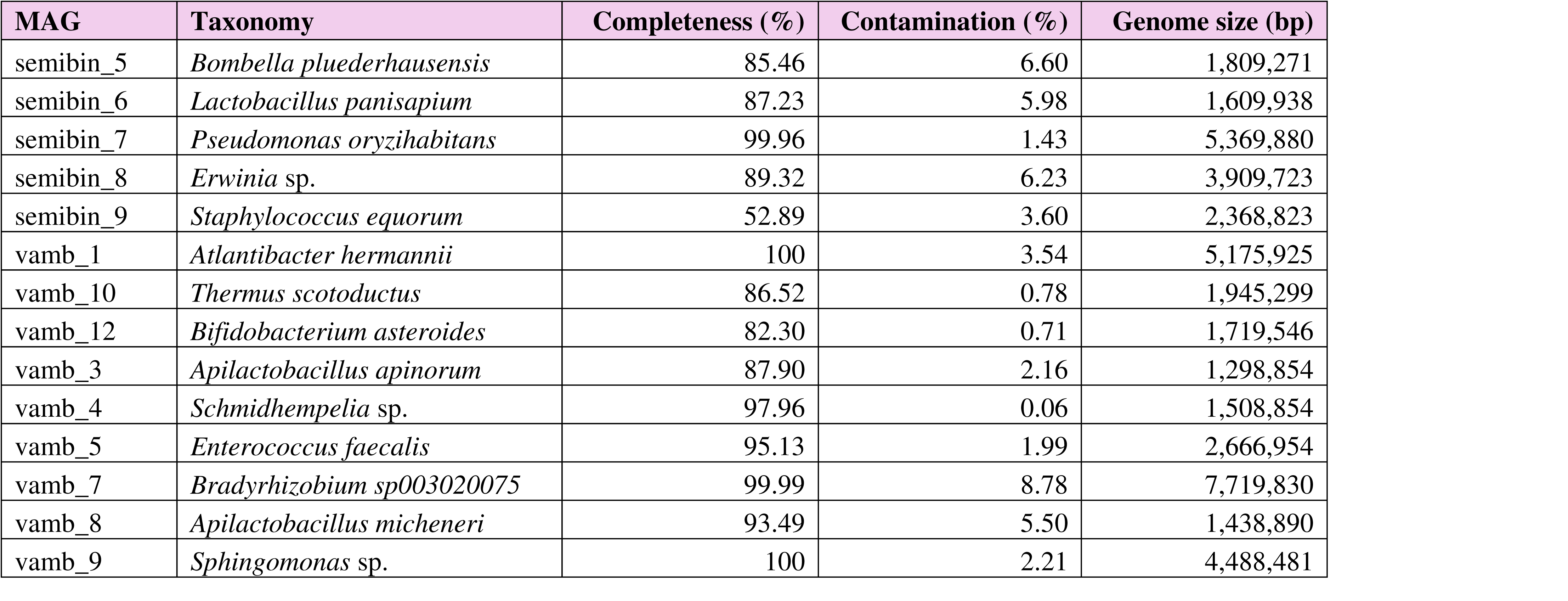
Taxonomic assignment and quality assessment of MAGs from metagenomic data.

### Identification and diversity of biosynthetic gene clusters

A total of 342 BGC sequences were identified, including 22 hybrid BGCs. When individual components within hybrid BGC sequences were counted separately (e.g., a hybrid NRPS + T1PKS cluster was counted as one NRPS-class BGC and one T1PKS-class BGC), the total number of BGCs increased to 366. The majority of the identified BGC sequences were located within MAGs (240/342).

The BGCs were classified into 38 classes of secondary metabolites. Among them, *terpene-precursor* and *terpene* were the two most abundant classes (**Figure 1a**), accounting for 16.7% and 15.8% of the BGCs, respectively. Besides, *RiPP-like* and *NRPS* groups were also detected in substantial numbers.

**Fig. 1.**
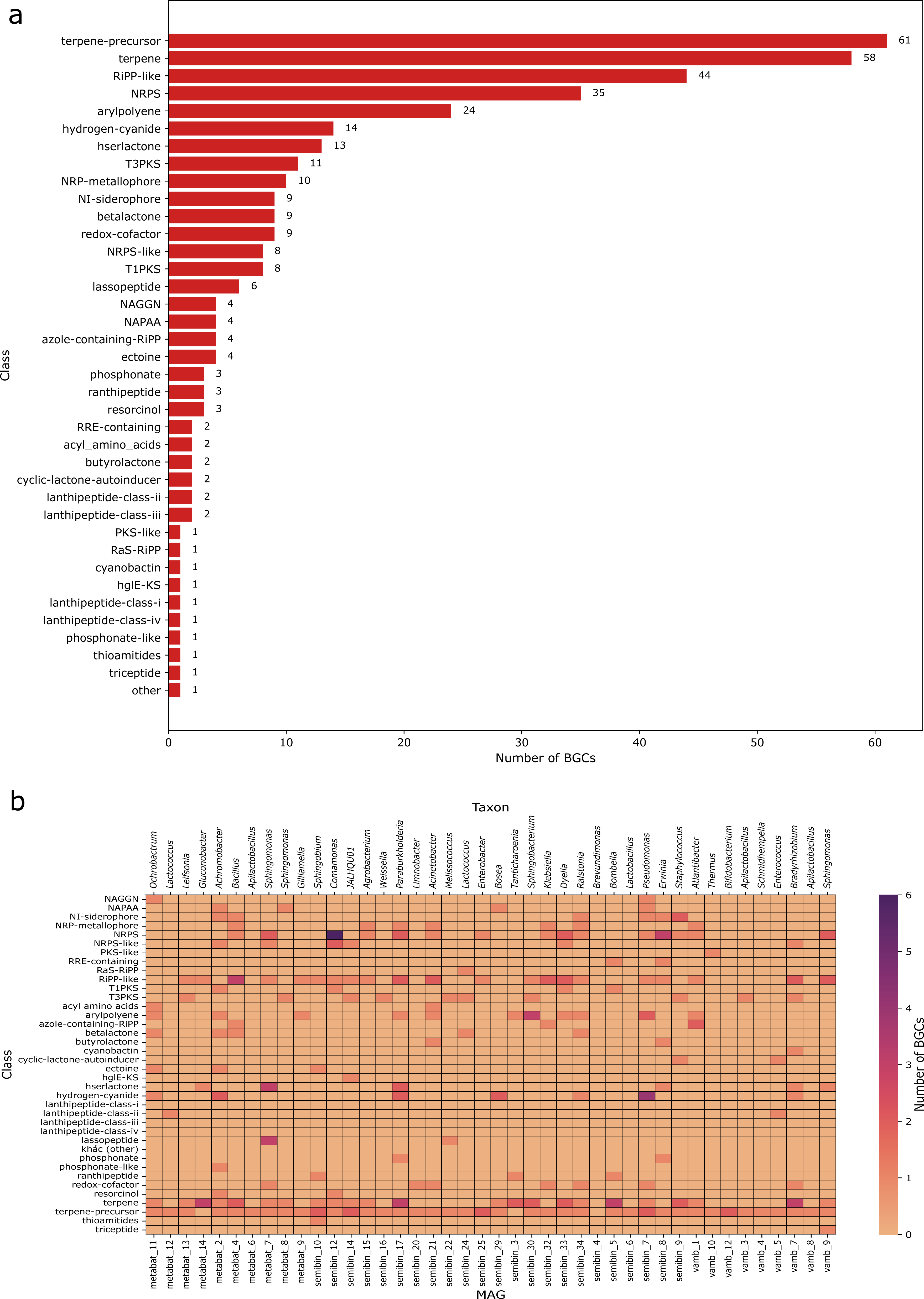
(a) Total number of BGCs by compound class; (b) The number of BGCs per MAG across classes

The heatmap (**Figure 1b**) illustrating the distribution of BGCs across compound classes revealed pronounced differences among MAGs. Most MAGs harbored only a limited number of BGCs, whereas a few MAGs represented biosynthetic hotspots, characterized by both a high number and diversity of BGCs. *Terpene* and *terpene-precursor* BGCs were present in the majority of MAGs.

### Similarity network analysis

The similarity network of BGCs comprised 31 gene cluster families (GCFs), including nine GCFs containing three or more BGCs and 22 doubletons (GCFs consisting of only two BGCs) (**Figure 2**). In addition, 276 singleton BGC sequences showed no similarity to any other BGCs in the dataset. Among the 31 identified GCFs, only 15 contained BGCs from the MIBiG database. Overall, 304 BGCs lacked corresponding reference sequences in MIBiG.

**Fig. 2.**
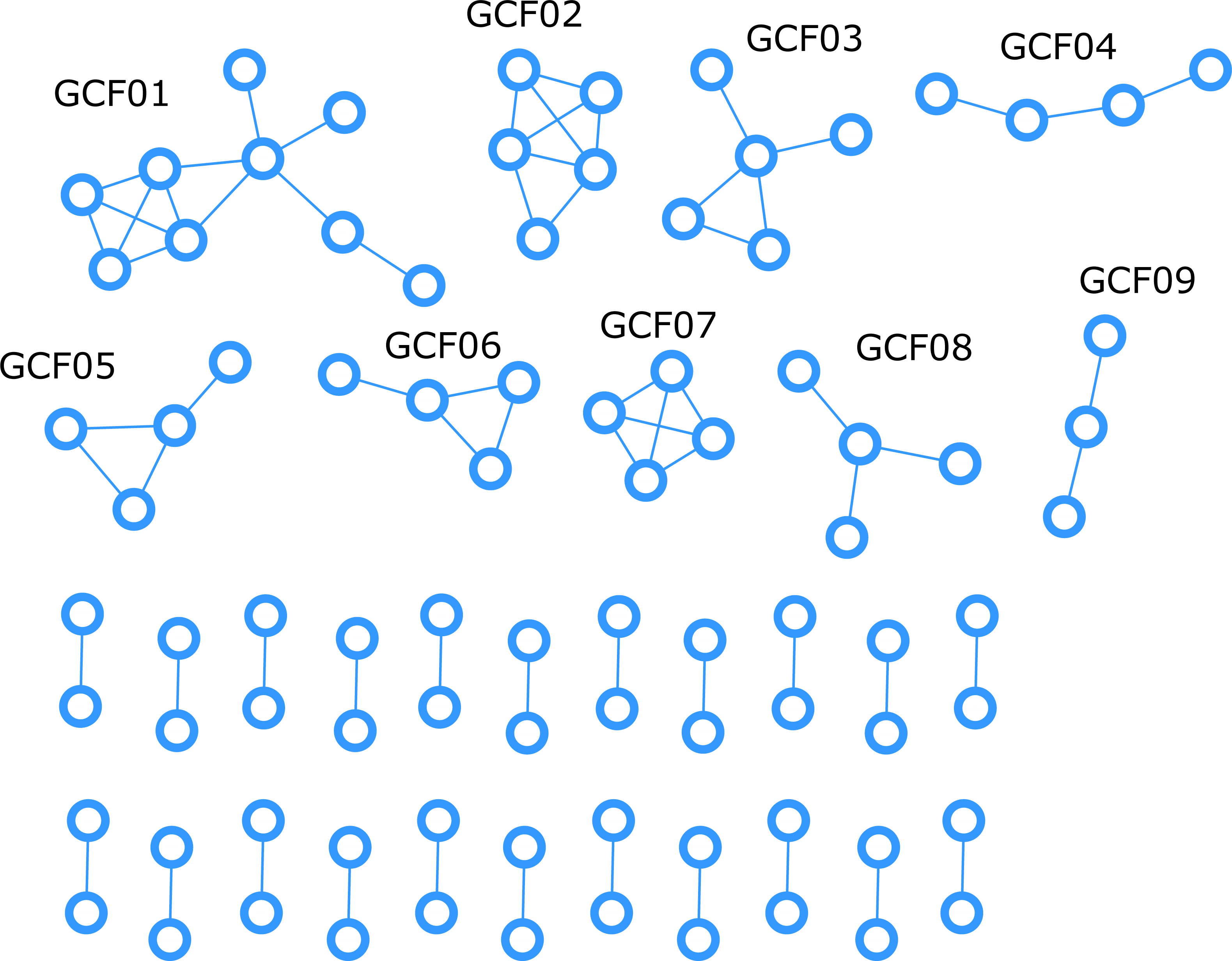
Similarity network of homologous BGC sequences. BGCs from the MIBiG database were included as references. Singletons were not shown

The nine GCFs containing three or more BGCs (named GCF01 to GCF09) were further analyzed in terms of their function. Among these, five GCFs contained BGCs that showed high similarity score to MIBiG entries (**Table 2**). Information derived from the reference BGCs indicated that GCF01, GCF03, and GCF08 are involved in siderophore biosynthesis. In addition, GCF04, GCF05, and GCF06 comprise aryl polyene–class BGCs. GCF02 and GCF09 are associated with terpene-precursor biosynthesis, whereas GCF07 is involved in the biosynthesis of azole-containing RiPPs. As the objective of this study was to explore novel bioactive compounds with antimicrobial potential, we selected GCF07 for further analysis. For clarity, BGCs are hereafter referred to by the names of the MAGs containing them (vamb 1, semibin 32, semibin 25, and unbinned).

**Table 2.**
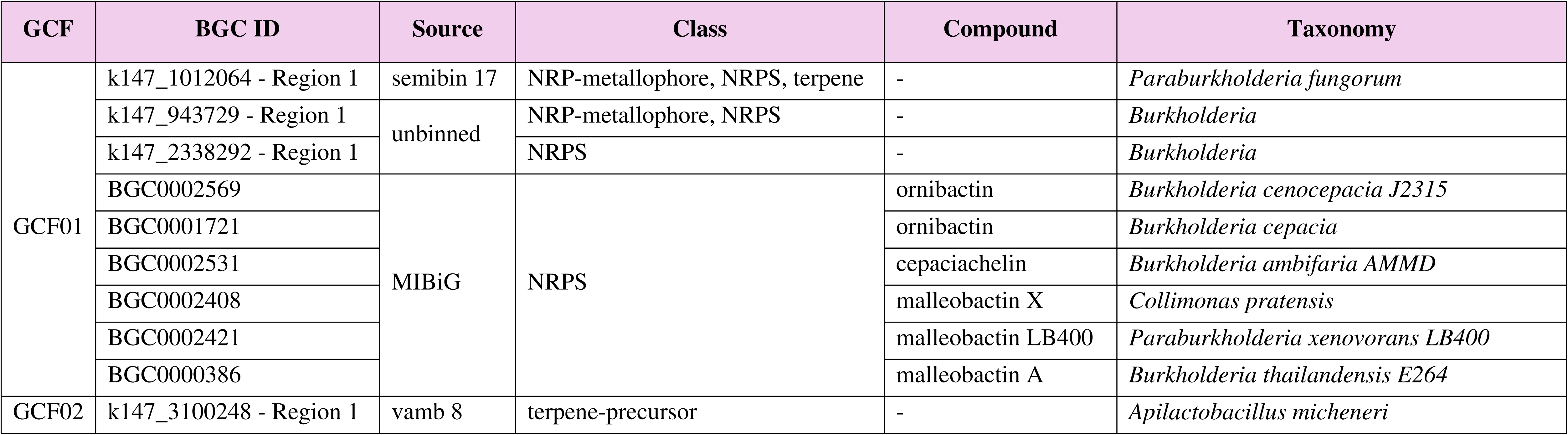
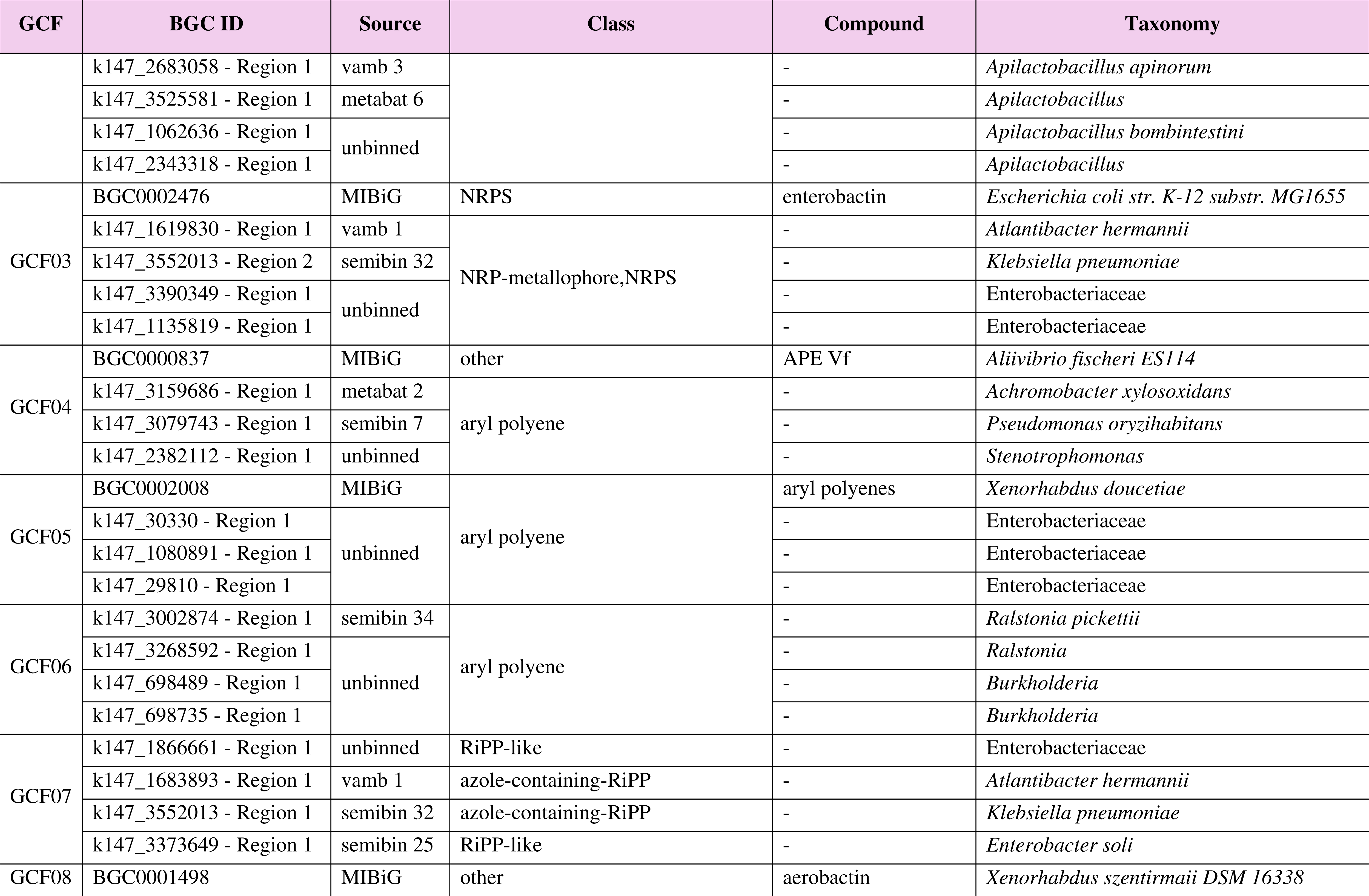
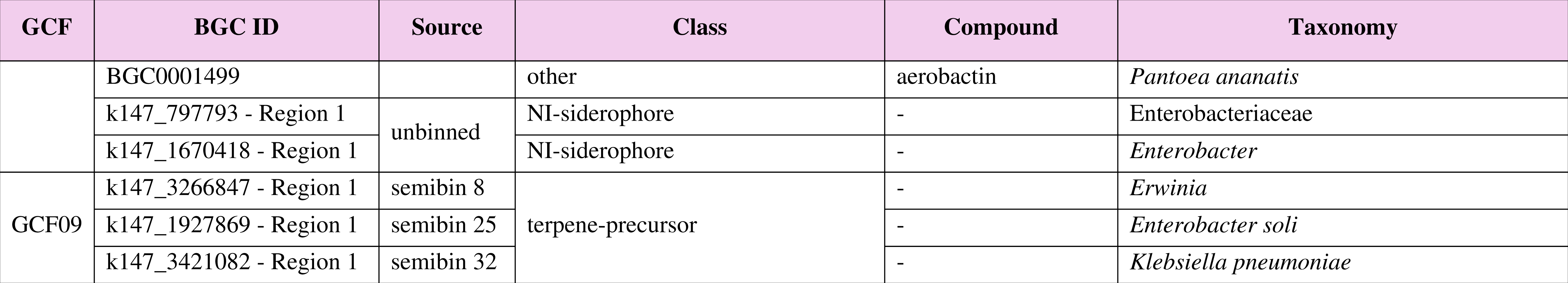
Information on GCFs. Only GCFs containing three or more BGCs are included.

| GCF | BGC ID | Source | Class | Compound | Taxonomy |
| --- | --- | --- | --- | --- | --- |
| GCF01 | k147_1012064 - Region 1 | semibin 17 | NRP-metallophore, NRPS, terpene | - | *Paraburkholderia fungorum* |
|  | k147_943729 - Region 1 | unbinned | NRP-metallophore, NRPS | - | *Burkholderia* |
|  | k147_2338292 - Region 1 |  | NRPS | - | *Burkholderia* |
|  | BGC0002569 | MIBiG | NRPS | ornibactin | *Burkholderia cenocepacia* J2315 |
|  | BGC0001721 |  |  | ornibactin | *Burkholderia cepacia* |
|  | BGC0002531 |  |  | cepaciachelin | *Burkholderia ambifaria* AMMD |
|  | BGC0002408 |  |  | malleobactin X | *Collimonas pratensis* |
|  | BGC0002421 |  |  | malleobactin LB400 | *Paraburkholderia xenovorans* LB400 |
|  | BGC0000386 |  |  | malleobactin A | *Burkholderia thailandensis* E264 |
| GCF02 | k147_3100248 - Region 1 | vamb 8 | terpene-precursor | - | *Apilactobacillus micheneri* |
|  | k147_2683058 - Region 1 | vamb 3 |  | - | *Apilactobacillus apinorum* |
|  | k147_3525581 - Region 1 | metabat 6 |  | - | *Apilactobacillus* |
|  | k147_1062636 - Region 1 | unbinned |  | - | *Apilactobacillus bombintestini* |
|  | k147_2343318 - Region 1 |  |  | - | *Apilactobacillus* |
| GCF03 | BGC0002476 | MIBiG | NRPS | enterobactin | *Escherichia coli* str. K-12 substr. MG1655 |
|  | k147_1619830 - Region 1 | vamb 1 | NRP-metallophore,NRPS | - | *Atlantibacter hermannii* |
|  | k147_3552013 - Region 2 | semibin 32 |  | - | *Klebsiella pneumoniae* |
|  | k147_3390349 - Region 1 | unbinned |  | - | Enterobacteriaceae |
|  | k147_1135819 - Region 1 |  |  | - | Enterobacteriaceae |
| GCF04 | BGC0000837 | MIBiG | other | APE Vf | *Aliivibrio fischeri* ES114 |
|  | k147_3159686 - Region 1 | metabat 2 | aryl polyene | - | *Achromobacter xylosoxidans* |
|  | k147_3079743 - Region 1 | semibin 7 |  | - | *Pseudomonas oryzihabitans* |
|  | k147_2382112 - Region 1 | unbinned |  | - | *Stenotrophomonas* |
| GCF05 | BGC0002008 | MIBiG | aryl polyene | aryl polyenes | *Xenorhabdus doucetiae* |
|  | k147_30330 - Region 1 | unbinned |  | - | Enterobacteriaceae |
|  | k147_1080891 - Region 1 |  |  | - | Enterobacteriaceae |
|  | k147_29810 - Region 1 |  |  | - | Enterobacteriaceae |
| GCF06 | k147_3002874 - Region 1 | semibin 34 | aryl polyene | - | *Ralstonia pickettii* |
|  | k147_3268592 - Region 1 | unbinned |  | - | *Ralstonia* |
|  | k147_698489 - Region 1 |  |  | - | *Burkholderia* |
|  | k147_698735 - Region 1 |  |  | - | *Burkholderia* |
| GCF07 | k147_1866661 - Region 1 | unbinned | RiPP-like | - | Enterobacteriaceae |
|  | k147_1683893 - Region 1 | vamb 1 | azole-containing-RiPP | - | *Atlantibacter hermannii* |
|  | k147_3552013 - Region 1 | semibin 32 | azole-containing-RiPP | - | *Klebsiella pneumoniae* |
|  | k147_3373649 - Region 1 | semibin 25 | RiPP-like | - | *Enterobacter soli* |
| GCF08 | BGC0001498 | MIBiG | other | aerobactin | *Xenorhabdus szentirmaii* DSM 16338 |
|  | BGC0001499 |  | other | aerobactin | *Pantoea ananatis* |
|  | k147_797793 - Region 1 | unbinned | NI-siderophore | - | Enterobacteriaceae |
|  | k147_1670418 - Region 1 |  | NI-siderophore | - | *Enterobacter* |
| GCF09 | k147_3266847 - Region 1 | semibin 8 | terpene-precursor | - | *Erwinia* |
|  | k147_1927869 - Region 1 | semibin 25 |  | - | *Enterobacter soli* |
|  | k147_3421082 - Region 1 | semibin 32 |  | - | *Klebsiella pneumoniae* |

The vamb 1 (*Atlantibacter hermannii*) cluster showed a high probability of producing compounds with antibacterial activity (74.5%). In contrast, semibin 32 (*Klebsiella pneumoniae*) displayed a relatively high antibacterial activity (50.8%), and also exhibited antifungal (33.5%) and notable antitumor activities (29.3%). Conversely, semibin 25 (*Enterobacter soli*) and the unbinned cluster did not show any significant predicted activities (**Table 3**). These two BGCs also contained only 3–5 genes, suggesting that they are likely incomplete BGCs (**Figure 3a**).

**Table 3.** Predicted proportions of biological activities of compounds biosynthesized by BGCs in GCF07.

| Bioactivity class | vamb 1 | semibin 25 | semibin 32 | unbinned |
| --- | --- | --- | --- | --- |
| antibacterial | 74,5% | 32,1% | 50,8% | 51,9% |
| antifungal | 19,9% | 3,0% | 33,5% | 21,9% |
| antitumor | 11,6% | 11,7% | 29,3% | 18,5% |
| surfactant | 34,7% | 17,7% | 7,6% | 1,5% |
| antiprotozoal | 0,5% | 0,1% | 0,4% | 0,1% |
| antiviral | 0,1% | 0,1% | 0,0% | 0,1% |
| siderophore | 1,6% | 7,7% | 0,2% | 0,6% |

The vamb 1 BGC contains a centrally located gene (**Figure 3b**) encoding a short peptide consisting of 56 amino acids (hereafter referred to as peptide V1). The amino acid sequence of peptide V1 is as follows: *MSALRKCRMVEGQPGETAYLTANMSVLLRQADRFCCVTAAAFYRRRVLAIADRVTE*

**Fig. 3.**
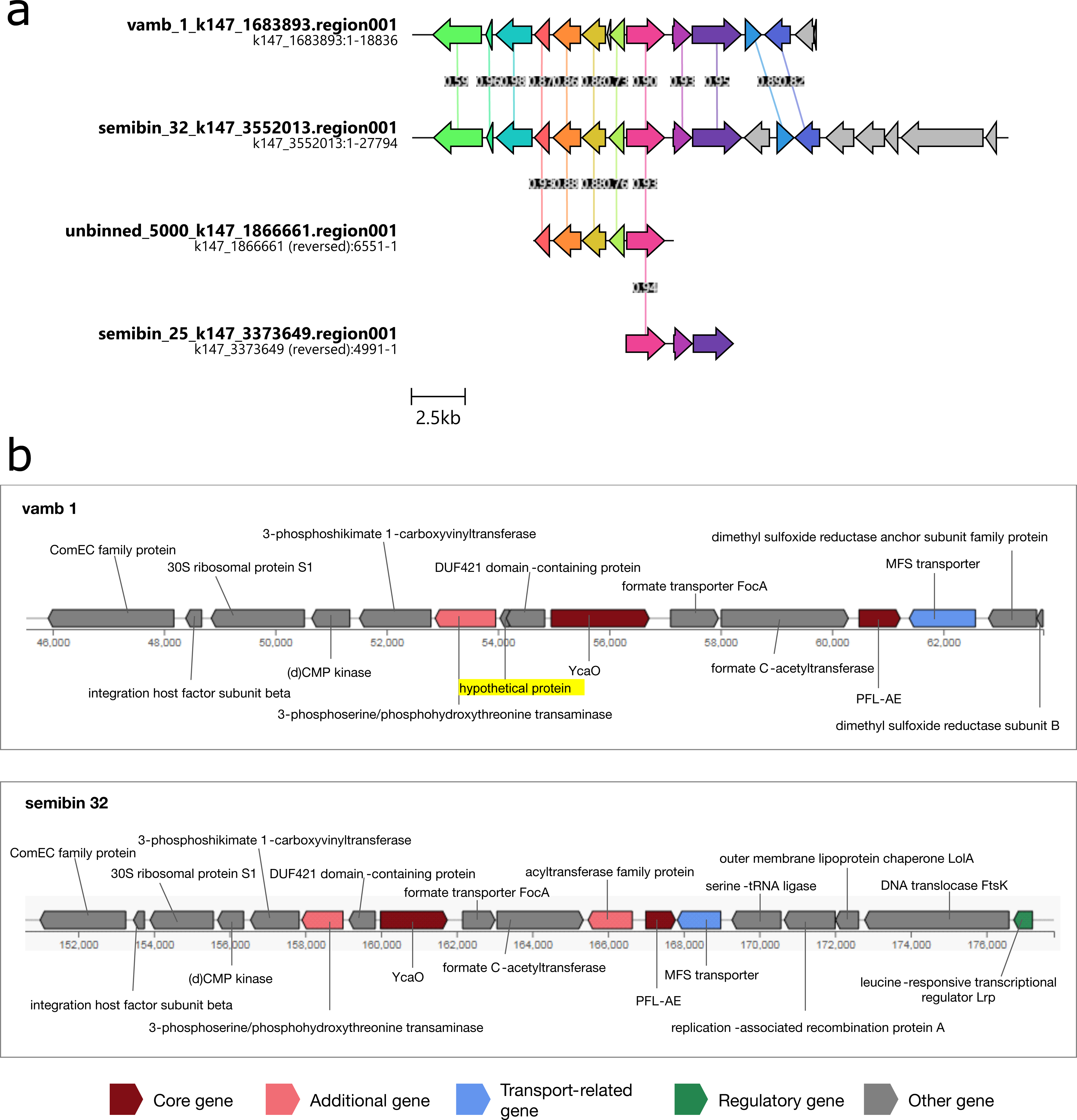
(a) Organization of BGCs within GCF07; (b) functional annotation of genes in the vamb 1 and semibin 32 BGCs. The vamb 1 cluster contains a short gene encoding a hypothetical protein (highlighted) that is proposed to be a RiPP precursor peptide

BLASTp analysis revealed that this peptide shares high sequence similarity (> 98%) with several hypothetical proteins from the genus *Klebsiella* in the NCBI *nr* database. These reference proteins have not yet been functionally characterized. However, both RiPPMiner and NeuRiPP predicted peptide V1 to be a precursor peptide of a RiPP-class compound.

## Discussion

### The honey microbiome represents a promising source of BGCs

This study provides evidence that the bacterial community associated with honey harbors substantial biosynthetic potential for the production of novel antimicrobial compounds. The identification of a large number of BGCs, most of which lack close matches in the MIBiG database, indicates that the honey microbiome represents an underexplored reservoir of secondary metabolite diversity. A large proportion of the BGCs identified in this study belong to the terpene and terpene-precursor classes. Terpenes constitute a major group of secondary metabolites with well-documented antioxidant and antibacterial activities, as well as therapeutic potential against various diseases, including cancer (Gonzalez-Burgos and Gomez-Serranillos 2012; Helfrich et al. 2019). In addition, the presence of RiPP and NRPS BGCs with prominent predicted bioactivity profiles further supports the notion that honey-associated bacteria may contribute to the discovery of new antimicrobial agents. These two classes encode ribosomally synthesized peptides and non-ribosomal peptides, respectively, which are well known as the sources of numerous clinically important natural antibiotics, such as nisin and penicillin (Awan et al. 2017; Zhao and Kuipers 2021).

### Metagenomics is an effective approach for exploring BGCs from honey-associated bacteria

Our results further demonstrate that metagenomics is a suitable and powerful approach for exploring BGC diversity in complex microbial communities. The recovery of MAGs and the detection of hundreds of BGCs, including many from uncultivated or poorly characterized taxa, highlight the advantage of metagenomic methods over culture-dependent strategies. In particular, network-based analyses enabled the identification of gene cluster families with limited or no representation in reference databases, thereby expanding the known biosynthetic landscape. Nevertheless, metagenomics primarily provides predictive insights and should be regarded as a hypothesis-generating tool that requires downstream experimental validation.

### A promising BGC candidate for new antimicrobial discovery

Among the identified gene cluster families, GCF07 emerged as a particularly promising candidate for further investigation due to its classification as an azole-containing RiPP and its enriched predicted antimicrobial activity. Azole-containing peptides have been widely reported to exhibit valuable biological activities, including antibacterial and antifungal effects (Emami et al. 2023). In this GCF, the vamb 1 BGC was prioritized based on its high predicted antibacterial potential and the presence of a short, centrally located precursor peptide. Independent predictions by RiPPMiner and NeuRiPP consistently support the role of this peptide as a RiPP precursor. Future studies should therefore focus on detailed characterization of the precursor peptide and associated tailoring enzymes, heterologous expression of the gene cluster, and comprehensive chemical and biological assays to confirm the structure and antimicrobial activity of the resulting compound.

### Limitations and future perspectives

Discovering novel secondary metabolites with new chemical structures using culture-based techniques remains challenging due to the frequent silencing of many BGCs under laboratory conditions. In this study, MAGs were recovered to enable the identification of BGCs directly from environmental sequencing data. Although this approach is widely used and practical, several limitations should be acknowledged. First, functional and bioactivity predictions are based entirely on *in silico* analyses and have not yet been experimentally validated. In addition, substantial portions of sequencing data may be discarded during the multiple filtering and assembly steps required for MAG reconstruction, potentially leading to the inadvertent loss of valuable sequences. Furthermore, metagenomic assembly frequently results in incomplete BGCs. The application of long-read sequencing technologies or increased sequencing depth may help to mitigate these limitations. Despite these challenges, this study establishes a robust framework for the systematic discovery of bioactive compounds from honey-associated microbial communities and provides a solid foundation for future experimental validation and exploration.

### Conclusions

This study provides a comprehensive metagenomic characterization of the biosynthetic potential of honey-associated bacteria from *Apis cerana* honey collected in the Northwest mountainous region of Vietnam. The identification of 366 BGCs spanning 38 compound classes, of which over 83% lack reference sequences in current databases, establishes this honey microbiome as a rich and largely untapped reservoir of biosynthetic novelty. The detection of an azole-containing RiPP BGC in a MAG assigned to *Atlantibacter hermannii*, predicted to exhibit strong antibacterial activity, provides a concrete and prioritized target for downstream experimental work, including heterologous expression and bioactivity validation. Collectively, these findings advance our understanding of the functional diversity of honey-associated microbial communities and lay the groundwork for future cultivation-based and biosynthetic approaches to access novel antimicrobial agents from underexplored environmental microbiomes.

## Declarations

### Author Contributions

All authors contributed to the conception and design of the study. Material preparation, data collection, and analysis were performed by Hoang-Nam Nguyen. Hoang-Nam Nguyen and Oanh T. P. Kim prepared the first draft of the manuscript. All authors edited and approved the manuscript.

